# Biodiversity on Indigenous lands equals that in protected areas

**DOI:** 10.1101/321935

**Authors:** Richard Schuster, Ryan R. Germain, Joseph R. Bennett, Nicholas J. Reo, Dave L. Secord, Peter Arcese

## Abstract

Declines in global biodiversity due to land conversion and habitat loss are driving a “Sixth Mass Extinction” and many countries currently fall short of meeting even nominal land protection targets to mitigate this crisis. Here, we quantify the potential contribution of Indigenous lands to biodiversity conservation using case studies of Australia, Brazil and Canada. Indigenous lands in each country are slightly more species rich than existing protected areas and, in Brazil and Canada, support more threatened species than existing protected areas or random sites. These results indicate that Indigenous lands and existing protected areas are similar in biodiversity. Enhanced partnerships between Indigenous communities and federal or state governments could help ameliorate current shortfalls in global biodiversity protection by facilitating protection for native species, helping to stem global biodiversity loss.

## Introduction

Habitat loss and degradation due to human land conversion are key threats to global biodiversity (Sala et al. 2000; Maxwell et al. 2016) that, to date, have been addressed mainly by expanding protected areas (PAs) globally. However, this approach has severe limitations, as many existing PAs have limited overlap with the geographic ranges of the world’s most threatened species (Rodrigues et al. 2004; Venter et al. 2014; Sánchez-Fernández & Abellán 2015), and in several regions of the world PAs overlap the ranges of endemic, high-priority species less often than expected if they had been located randomly (Nori et al. 2015; Sánchez-Fernández & Abellán 2015). Thus, even given a rapid increase in the global extent of PAs to meet the Convention on Biological Diversity’s (CBD) target of protecting 17% of global terrestrial area, shortfalls in coverage and implementation suggest 17% will be insufficient to prevent further extinctions or other coverage-based conservation goals (Watson et al. 2014; Barnes 2015; Polak et al. 2016). This implies that many species are destined to extinction unless they can maintain positive growth rates on land not being set aside as PAs, such as Indigenous and other community-managed lands.

Indigenous land tenure practices have in many cases led to higher native and rare species richness (Redford & Stearman 1993; Peres 1994; Yibarbuk et al. 2001; Arcese et al. 2014) and less deforestation and land degradation than non-indigenous practices (Waller & Reo, in press; Nepstad et al. 2006; Porter-Bolland et al. 2012; Nolte et al. 2013; Ceddia et al. 2015). However, despite these indicators that Indigenous lands (i.e., land parcels managed or co-managed by Indigenous communities) could play a large positive role in biodiversity conservation, no study has quantified patterns of the biodiversity or conservation value of Indigenous lands in separate global regions, or estimated their contribution to global biodiversity conservation in the future. We show how Indigenous lands can help to meet international biodiversity conservation targets, and discuss governance arrangements that could protect and expand the role Indigenous peoples play in supporting biodiversity conservation globally.

Specifically, we quantified the contributions of Indigenous lands to biodiversity in three countries (Australia, Brazil, Canada) with large total areas owned and/or managed by Indigenous communities. To do so, we estimated total richness of amphibians, birds, mammals and reptiles using freely available data from International Union for the Conservation of Nature (IUCN) species range maps. We then compared estimates among three land types: Indigenous lands, protected areas and randomly selected sites of equivalent area. We similarly estimated the richness of species at risk (i.e., classified as ‘vulnerable’, ‘endangered’, and ‘critically endangered’ by the IUCN). Maps of PAs were extracted from the World Database on Protected Areas (WDPA) and defined as IUCN designated PA categories I–VI. Delineations of Indigenous lands were extracted from WDPA, Australian Land Tenure and the Collaborative Australian Protected Areas Database for Australia, from WDPA for Brazil, and from National Resources Canada (Fig. 1, see Methods for details).

**Fig. 1.**
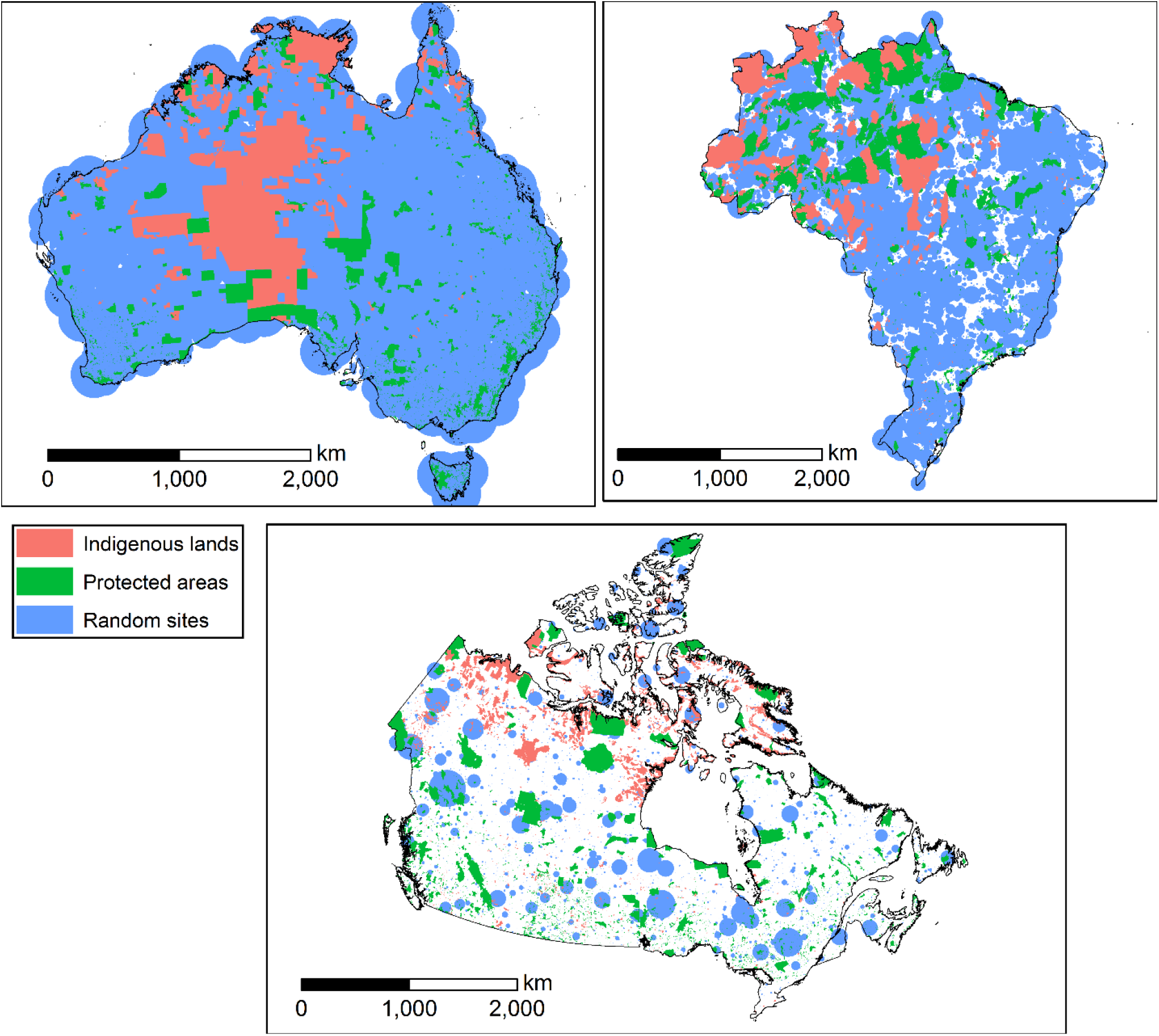
Location of Indigenous Lands, Protected Areas and Random Sites in Australia, Brazil and Canada.

## Methods

### Data processing

We based our analysis on an initial set of 26,688 spatial data layers collected from different sources and in different formats. This data set consisted of three basic administrative delineations (country boundaries, protected areas and Indigenous lands), and 26,682 species distributions, as described below, with total terrestrial land restricted to the country boundaries of Australia, Brazil and Canada.

### Basic administrative delineations

National boundaries were derived from the Global Administrative Areas database (http://gadm.org/, accessed 2015–10-10). The data on protected areas (PA) was based on the September 2016 release of the World Database on Protected Area (WDPA, https://www.protectedplanet.net). We extracted the protected areas for each country from the WDPA database by selecting only areas belonging to IUCN protected area categories I–VI and having as status ‘designated’. This resulted in totals of 7740, 1117 and 6764 protected areas for Australia, Brazil and Canada respectively. As there is to our knowledge no comprehensive database for Indigenous lands (IL) globally, we created country-specific Indigenous lands layers from several sources. For Australia we used WDPA, Australian Land Tenure (ALT) (Australian Government 2016a) and Collaborative Australian Protected Areas Database (CAPAD) (Australian Government 2014) data, totalling 308 polygons. From the WDPA data we included polygons which names started with ‘Aborig’ or ‘Indige’ for the Indigenous lands dataset and excluded them from the protected areas dataset; from ALT we included polygons with the RES_TYPE FINP (Aboriginal freehold national park), AA (Aboriginal Area), ABOR (Aboriginal Reserve), AFI (Aboriginal freehold land [inalienable]), AFL (Aboriginal freehold land [alienable]), ALL (Aboriginal held lease other than pastoral), ALG (Aboriginal local government area lease), AP (Aboriginal place [designated under New South Wales Wildlife Act]), APL (Aboriginal held pastoral lease), and AS (Aboriginal site); and from CAPAD data we included polygons with the types ‘Aboriginal Area’, ‘Indigenous Protected Area’ and ‘National Park Aboriginal’. For Brazil, the WDPA specifically delineates Indigenous lands; we utilized the following designations to select polygons: ‘Indigenous Area’ and ‘Indigenous Reserve’, totalling 718 polygons. For Canada, the federal government provides a ‘Canada Aboriginal Lands’ layer (Natural Resources Canada 2014), which we used here, totalling 3124 polygons. To compare species richness on protected areas and Indigenous lands to a benchmark we created 10,000 randomly located points in each of the three countries. We chose this number to create random sites in the same order of magnitude as there are PA and IL. These random points served as the centroids of circular ‘random areas’ (RA). The size of each of these areas was determined by creating a list of the sizes of protected areas and Indigenous lands and randomly assigning their sizes to each centroid. This way we ensured the creation of random areas comparable in size to the protected areas and Indigenous lands we investigated in each of the three countries, which themselves differ in size and shape (Fig. 1).

### Species

Our species lists were determined using the IUCN Red List of threatened species, following Pouzols et al. (2014). For mammal, amphibian and reptile species ranges, we used the IUCN Red List website (http://www.iucnredlist.org/, accessed 2016–09-14) and for birds we used the BirdLife International data zone webpage (http://www.birdlife.org/datazone/home, accessed 2016–09-14). These data have certain limitations, including possible underestimation of the extent of occurrence and overestimation of the true area of occupancy (Pouzols et al. 2014), but have been shown to be robust to commission errors as long as the focus is on species assemblages rather than single species, (Venter et al. 2014). They are currently the most frequently used and updated information on vertebrate species distributions globally (Le Saout et al. 2013).

For each species group, we restricted our analysis to species that fell into the presence category of ‘Extant’, the origin categories of ‘Native’ or ‘Reintroduced’ and the seasonality categories ‘Resident’, ‘Breeding Season’ or ‘Non-breeding Season’. For each country, we first selected each IUCN polygon that intersected its national border and subsequently we clipped each polygon by that border. This resulted in the following final numbers of amphibian, bird, mammal and reptile species per country: 219, 736, 270, 178 (Australia); 909, 1748, 621, 160 (Brazil); 50, 443, 157, 41 (Canada). In addition to analysing all species, we also analyzed threatened species only. All species with an IUCN status of ‘critically endangered’, ‘endangered’ and ‘vulnerable’ were assigned to the threatened category. This resulted in the following final numbers of threatened species of amphibians, birds, mammals and reptiles per country: 48, 57, 53, 17 (Australia); 36, 159, 78, 20 (Brazil); 1, 14, 6, 9 (Canada).

### Analysis

The analysis steps for each species in each of the three countries were identical, and consisted of first creating a shapefile for each of the 26682 species distributions from the combined shapefiles or geodatabases for each species group. For ease of processing we then split each of the species polygons up into smaller segments of a maximum size of 250 × 250km. Subsequently we intersected each species shapefile with IL, PA and RA and retained the areas of overlap (QGIS). We then summarized results and calculated generalized linear regression models (Negative binomial, link = log) comparing IL, PA and RA for total species richness, species richness by species group, total threatened species richness and threatened species richness by species group. We further created density plots to visualize the relationship the relationship between species richness and IL, PA and RA. Analyses were conducted in R (R Development Core Team 2016). Data, scripts and full results are available here: https://osf.io/f86wv/?view_only=a963ae92f68041ad97fb3ac3ccc4616d

## Results

Indigenous lands, as legally recognized by the three national governments, represent 16.5, 13.3, and 6.3% of terrestrial area for Australia, Brazil and Canada, respectively. PAs represent 9.2, 21.1, and 10.7% of terrestrial area for Australia, Brazil and Canada, respectively. In all three countries, Indigenous lands have the highest species richness in all focal taxonomic groups combined, with randomly selected areas having the lowest species richness (Fig. 2 a,b,c). Indigenous lands also have higher species richness than randomly selected areas for each focal taxonomic group for all three countries (Appendix S1, S2), and slightly higher species richness than protected areas (PAs) for all focal taxonomic groups in Brazil, for all groups except birds in Australia, and for mammals and amphibians in Canada (Appendix S1, S2). Threatened species richness of all taxa combined was also higher on Indigenous lands than randomly selected areas for all three countries, and slightly higher than in PAs for Brazil and Canada (Table 1). In addition, threatened species richness was higher on Indigenous lands than PAs for amphibians and reptiles in Australia, mammals in Brazil, and birds and reptiles in Canada (Appendix S3).

**Fig. 2.**
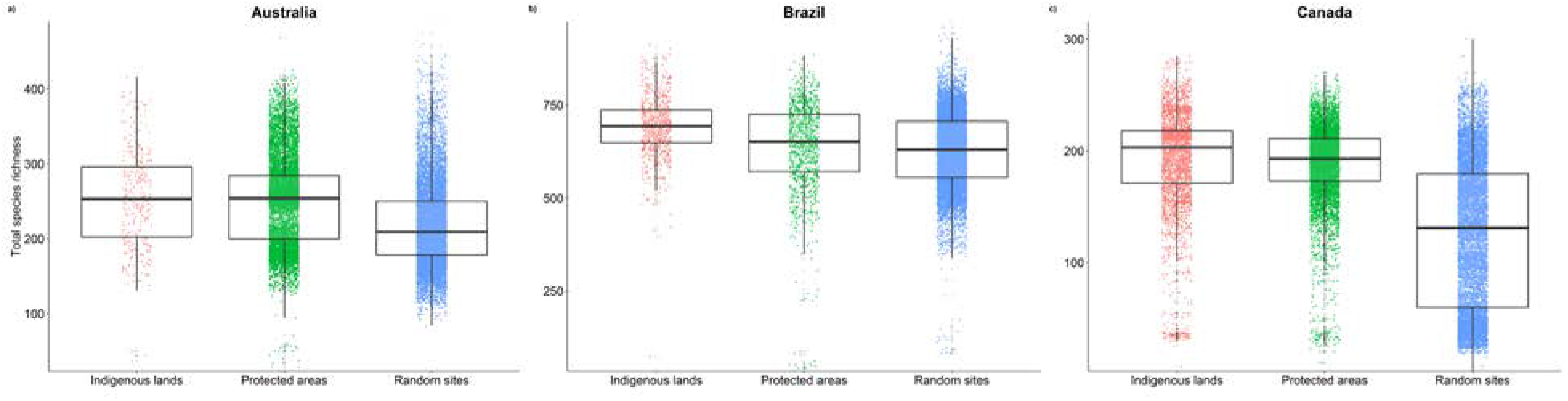
Total species richness for a) Australia, b) Brazil, c) Canada on Indigenous lands, protected areas and random sites. Colored jitter plots show the distribution of the raw data and the boxplots show summarized data in form median, first and third quartile.

**Table 1.** Summaries of country-specific linear regressions comparing species richness. Each model utilizes a negative binomial error structure and compares species richness on Indigenous lands, protected areas and random sites. 95% confidence interval values are presented in parenthesis. In all cases but one (red listed species in Australia), IL outperforms protected areas. IL further outperforms random sites all models tested.

| **All Species** | **Indigenous Lands** | **Protected Areas** | **Random sites** |
| --- | --- | --- | --- |
| Australia | 253 (246 - 260) | 250 (249 - 252) | 218 (217 - 219) |
| Brazil | 692 (683 - 701) | 639 (633 - 645) | 628 (626 - 630) |
| Canada | 192 (189 - 195) | 189 (187 - 191) | 126 (125 - 127) |
| **Red Listed Species** |  |  |  |
| Australia | 5.86 (5.46 - 6.28) | 7.31 (7.22 - 7.4) | 4.19 (4.13 - 4.25) |
| Brazil | 15.88 (15.37 - 16.4) | 14.68 (14.3 - 15.07) | 15.21 (15.08 - 15.35) |
| Canada | 2.4 (2.34 - 2.46) | 2.01 (1.98 - 2.04) | 2.37 (2.34 - 2.4) |

To rule out potential confounding factors on the results presented in Fig. 2 and speculate on the potential mechanisms underlying the patterns of species richness across the three countries investigated, we further evaluated the effects of land patch size and geographic location (lat/long) on species richness in lands identified as Indigenous lands, protected areas, and randomly placed areas (Appendix S4). These analyses revealed no consistent patterns in terms of the effects of patch size or location on total species richness or the richness of species at risk across the three countries investigated, meaning that the consistent results across each country presented in Fig 2 are unlikely to be influenced by these potential confounding factors. Instead, we interpret our results to indicate that Indigenous community land tenure practices may themselves result in higher species richness than random land areas and roughly equivalent species richness to protected areas.

## Discussion

Our results indicate that Indigenous lands represent an important source of species richness in three separate countries, and could thus play a pivotal role in biodiversity conservation. We found that the distributions of more species, and more threatened species in particular, overlapped Indigenous lands than overlapped existing protected areas or randomly selected sites with all investigated taxa combined. Although differences in richness between Indigenous lands and protected areas were relatively small, our results imply that Indigenous lands and protected areas provide complimentary benefits to global conservation initiatives. Indeed, many species are largely dependent on Indigenous lands for their continued persistence. In Australia, the ranges of two threatened species (Scanty frog [*Cophixalus exiguous*], Northern hopping-mouse [*Notomys aquilo*]) total less than 5000km^2^, but of these areas > 97% is on Indigenous lands. Of all species considered, 42 in Australia, 216 in Brazil, and two species in Canada had > 50% of their range in a focal country on Indigenous lands, while 11 species at risk in Australia, 10 in Brazil, and zero in Canada had > 50% of their range in a focal country on Indigenous lands.

While PAs are a key tool in biodiversity conservation (Watson et al. 2014), Indigenous land management practices, as well as co-management between Indigenous communities and federal or state governments, have also been shown to contribute positively to biodiversity (Gilligan 2006; Ricketts et al. 2010) and our results indicate that Indigenous stewardship may be essential to the persistence of some species in the future given the high degree of overlap between the ranges of threatened species and Indigenous lands. Independent, Indigenous land tenure as well as collaborative approaches to conservation linking Indigenous and non-Indigenous institutions appear necessary to achieve CBD goals in the future for Australia, Brazil and Canada. Such mutually-beneficial partnerships have been demonstrated at the local scale, in terms of the contributions of Indigenous lands to the mature forests and sustainable wildlife populations on non-Indigenous forestlands (Waller & Reo, in press). Larger-scale collaborative approaches to conservation are used across the globe because of their potential to address underlying causes of biodiversity loss that individual governments and organizations cannot address on their own (Sayer et al. 2013). Although successful partnerships between environmental 183s and Indigenous communities have historically been relatively rare (LaDuke 2017), empirical studies have revealed some guiding principles for burgeoning partnerships. For instance, Stevens and colleagues (Stevens 1997) created a series of principles designed to guide the formation of a new paradigm for parks and PAs that fully embraces Indigenous rights and land tenure systems. Further studies have characterized commonalities in successful partnerships between Indigenous and non-Indigenous groups in terms of achieving environmental goals and determined that such partnerships are constituted in ways that respect Indigenous nations’ political and governmental authority and cultural distinctiveness (Reo et al. 2017, Reo et al. in press).

The establishment of national parks and protected areas has had severe social consequences for Indigenous peoples around the world since the early 1900’s (Stevens 1997; West et al. 2006). Protected areas established based on Euro-American wilderness ideals prohibited Indigenous peoples from exercising their traditional ways of life and forcibly removed them from their homelands. Removal of Indigenous peoples from their homelands led to negative consequences for the Indigenous societies (West et al. 2006) and often for the ecosystems that conservationists aimed to “protect” (Gomez-Pompa & Kaus 1992; Stevens 1997). Our results suggest that collaboration between Indigenous and non-Indigenous land managers could result in synergies that dramatically advance conservation. Models for such partnerships might include providing resources to facilitate stewardship and implementation. For instance, the Canadian federal parks agency spends *c* $ 282 per km^2^ ($ 92M CAD/yr) on programs related to the enforcement, restoration and maintenance of biodiversity in protected areas (Parks Canada 2016). Similar expenditures to subsidize traditional management practices on Indigenous lands in Canada would imply a cost of $ 176M CAD/yr based on costs per unit areas. Mutually agreed co-management arrangements offer an additional route to collaboration between federal governments and Indigenous landowners or managers. Co-management is already common in Australia, where Indigenous Protected Areas (IPAs) occur wholly on Indigenous lands and make up > 40% of the national protected area system (Australian Government 2016b). Some IPAs are co-managed with non-Indigenous partners as part of land claims agreements, while others are under sole Indigenous management with funding and technical assistance from non-Indigenous governments (Bauman & Smyth 2007; Ross et al. 2009).

Although many countries are on track to meet the nominal 17% terrestrial protected area goal of the Convention on Biological Diversity (Butchart et al. 2015), meeting the goals for representativeness and connectivity of protected areas or other effective area-based conservation measures will be challenging. Meeting the CBD’s target of preventing extinction is likely to require much larger protected area networks than anticipated in the area-based goal (Barnes 2015; Polak et al. 2016). Recognizing the role of Indigenous lands and leadership in biodiversity conservation, and facilitating voluntary partnerships to ensure the conservation of habitats on Indigenous lands, may provide crucial opportunities for many countries to meet their international commitments to biodiversity conservation, while supporting traditional land management.

## Acknowledgments

We thank N. Baron, A. Jacob, H. Locke, M.-C. Loretto, A. Rodewald, and O. Venter for providing comments on the manuscript. RS is supported by a Liber Ero Fellowship, and RRG, JRB, and PA are supported by the Natural Sciences and Engineering Research Council of Canada. The authors declare no competing interests. The data reported in this paper are tabulated in the Supplementary Materials. Data, analysis scripts and full results are archived here: https://osf.io/f86wv/?view_only=a963ae92f68041ad97fb3ac3ccc4616d.

## Supporting Information

Additional Supporting Information may be found in the online version of this article:

**S1.** This supplementary material provides detailed results of country-specific linear regressions for all species.

**S2.** This supplementary material provides taxa-specific density plots of species richness for each of the three countries investigated.

**S3.** This supplementary material provides detailed results of country-specific linear regressions for IUCN threatened species.

**S4.** This supplementary material provides detailed results of country-specific linear regressions for all species using a range of covariates to explain species richness.

## Author contributions

RS, RRG, and PA conceived the study. RS and RRG collected data and conducted analyses. All authors contributed to writing and editing the paper.

## References

Arcese, P., Schuster, R., Campbell, L., Barber, A. & Martin, T.G. (2014). Deer density and plant palatability predict shrub cover, richness, diversity and aboriginal food value in a North American archipelago. Divers. Distrib., 20, 1368–1378.

Australian Government. (2014). Collaborative Australian Protected Areas Database (CAPAD) 2014 [WWW Document]. URL http://www.environment.gov.au/fed/catalog/search/resource/details.page?uuid=%7B4448C ACD-9DA8–43D1-A48F-48149FD5FCFD% 7D

Australian Government. (2016a). Australian Land Tenure 1993 [WWW Document]. URL https://data.gov.au/dataset/australian-land-tenure-1993

Australian Government. (2016b). Indigenous Protected Areas – IPAS [WWW Document]. URL https://www.dpmc.gov.au/indigenous-affairs/environment/indigenous-protected-areas-ipas

Barnes, M. (2015). Aichi targets: Protect biodiversity, not just area. Nature, 526, 195.

Bauman, T. & Smyth, D. (2007). Indigenous partnerships in protected area management in Australia: three case studies. Australian Institute of Aboriginal and Torres Strait Islander Studies Canberra.

Butchart, S.H.M., Clarke, M., Smith, R.J., Sykes, R.E., Scharlemann, J.P.W., Harfoot, M., Buchanan, G.M., Angulo, A., Balmford, A., Bertzky, B., Brooks, T.M., Carpenter, K.E., Comeros-Raynal, M.T., Cornell, J., Ficetola, G.F., Fishpool, L.D.C., Fuller, R.A., Geldmann, J., Harwell, H., Hilton-Taylor, C., Hoffmann, M., Joolia, A., Joppa, L., Kingston, N., May, I., Milam, A., Polidoro, B., Ralph, G., Richman, N., Rondinini, C., Segan, D.B., Skolnik, B., Spalding, M.D., Stuart, S.N., Symes, A., Taylor, J., Visconti, P., Watson, J.E.M., Wood, L. & Burgess, N.D. (2015). Shortfalls and Solutions for Meeting National and Global Conservation Area Targets. Conserv. Lett., 8, 329–337.

Ceddia, M.G., Gunter, U. & Corriveau-Bourque, A. (2015). Land tenure and agricultural expansion in Latin America: The role of Indigenous Peoples’ and local communities’ forest rights. Glob. Environ. Chang., 35, 316–322.

Gilligan, B. (2006). The indigenous protected areas programme: 2006 evaluation. Department of the Environment and Heritage.

Gomez-Pompa, A. & Kaus, A. (1992). Taming the Wilderness Myth. Bioscience, 42, 271–279.

LaDuke, W. (2017). All our relations: Native struggles for land and life. Haymarket Books.

Maxwell, S.L., Fuller, R.A., Brooks, T.M. & Watson, J.E.M. (2016). Biodiversity: The ravages of guns, nets and bulldozers. Nature, 536, 143–145.

Natural Resources Canada. (2014). Aboriginal Lands of Canada [WWW Document]. URL http://geogratis.gc.ca/api/en/nrcan-rncan/ess-sst/815dd99d-4fbd-47cc-be02–7ad4b03a23ec.html#distribution

Nepstad, D., Schwartzman, S., Bamberger, B., Santilli, M., Ray, D., Schlesinger, P., Lefebvre, P., Alencar, A., Prinz, E., Fiske, G., Rolla, A. (2006). Inhibition of Amazon deforestation and fire by parks and indigenous lands. Conserv. Biol., 20, 65–73.

Nolte, C., Agrawal, A., Silvius, K.M. & Soares-Filho, B.S. (2013). Governance regime and location influence avoided deforestation success of protected areas in the Brazilian Amazon. Proc. Natl. Acad. Sci., 110, 4956–4961.

Nori, J., Lemes, P., Urbina-Cardona, N., Baldo, D., Lescano, J. & Loyola, R. (2015). Amphibian conservation, land-use changes and protected areas: A global overview. Biol. Conserv., 191, 367–374.

Parks Canada. (2016). 2016–17 Report on Plans and Priorities [WWW Document]. URL http://www.pc.gc.ca/eng/docs/pc/plans/rpp/rpp2016–17/index.aspx

Peres, C.A. (1994). Indigenous reserves and nature conservation in Amazonian forests. Conserv.Biol., 8, 586–588.

Polak, T., Watson, J.E.M., Bennett, J.R., Possingham, H.P., Fuller, R.A. & Carwardine, J. (2016). Balancing Ecosystem and Threatened Species Representation in Protected Areas and Implications for Nations Achieving Global Conservation Goals. Conserv. Lett.

Porter-Bolland, L., Ellis, E.A., Guariguata, M.R., Ruiz-Mallén, I., Negrete-Yankelevich, S. & Reyes-García, V. (2012). Community managed forests and forest protected areas: an assessment of their conservation effectiveness across the tropics. For. Ecol. Manag., 268, 6–17.

Pouzols, F.M., Toivonen, T., Di Minin, E., Kukkala, A.S., Kullberg, P., Kuusterä, J., Lehtomäki, J., Tenkanen, H., Verburg, P.H. & Moilanen, A. (2014). Global protected area expansion is compromised by projected land-use and parochialism. Nature, 516, 383–386.

R Development Core Team. (2016). R: A language and environment for statistical computing 3.3.2, http://www.r-project.org.

Redford, K.H. & Stearman, A.M. (1993). Forest-Dwelling Native Amazonians and the Conservation of Biodiversity: Interests in Common or in Collision? Conserv. Biol., 7, 248–

Reo, N.., Whyte, K.P., McGregor, D., Smith, M.A. & Jenkins, J. (In Press). Factors that support Indigenous involvement in multi-actor environmental stewardship. AlterNative.

Ricketts, T.H., Soares-Filho, B., da Fonseca, G.A.B., Nepstad, D., Pfaff, A., Petsonk, A., Anderson, A., Boucher, D., Cattaneo, A. & Conte, M. (2010). Indigenous lands, protected areas, and slowing climate change. PLoS Biol, 8, e1000331.

Rodrigues, A.S.L., Andelman, S.J., Bakarr, M.I., Boitani, L., Brooks, T.M., Cowling, R.M., Fishpool, L.D.C., Da Fonseca, G.A.B., Gaston, K.J. & Hoffmann, M. (2004). Effectiveness of the global protected area network in representing species diversity. Nature, 428, 640– 643.

Ross, H., Grant, C., Robinson, C.J., Izurieta, A., Smyth, D. & Rist, P. (2009). Co-management and Indigenous protected areas in Australia: achievements and ways forward. Australas. J. Environ. Manag., 16, 242–252.

Sala, O.E., Stuart Chapin, F., III, Armesto, J.J., Berlow, E., Bloomfield, J., Dirzo, R., Huber-Sanwald, E., Huenneke, L.F., Jackson, R.B., Kinzig, A., Leemans, R., Lodge, D.M., Mooney, H.A., Oesterheld, M., Poff, N.L., Sykes, M.T., Walker, B.H., Walker, M. & Wall, D.H. (2000). Global Biodiversity Scenarios for the Year 2100. Science (80–.)., 287, 1770–LP-1774.

Sánchez-Fernández, D. & Abellán, P. (2015). Using null models to identify under-represented species in protected areas: A case study using European amphibians and reptiles. Biol. Conserv., 184, 290–299.

Le Saout, S., Hoffmann, M., Shi, Y., Hughes, A., Bernard, C., Brooks, T.M., Bertzky, B., Butchart, S.H.M., Stuart, S.N., Badman, T. & others. (2013). Protected areas and effective biodiversity conservation. Science (80–.)., 342, 803–805.

Sayer, J., Sunderland, T., Ghazoul, J., Pfund, J.-L., Sheil, D., Meijaard, E., Venter, M., Boedhihartono, A.K., Day, M., Garcia, C., van Oosten, C. & Buck, L.E. (2013). Ten principles for a landscape approach to reconciling agriculture, conservation, and other competing land uses. Proc. Natl. Acad. Sci. U. S. A., 110, 8349–56.

Stevens, S. (1997). Conservation through cultural survival: Indigenous peoples and protected areas. Island Press.

Venter, O., Fuller, R.A., Segan, D.B., Carwardine, J., Brooks, T., Butchart, S.H.M., Di Marco, M., Iwamura, T., Joseph, L., O’Grady, D., Possingham, H.P., Rondinini, C., Smith, R.J., Venter, M. & Watson, J.E.M. (2014). Targeting Global Protected Area Expansion for Imperiled Biodiversity. PLOS Biol., 12, e1001891.

Waller, D.M. & Reo, N.J. (In Press). First Stewards: Ecological outcomes of forest and wildlife stewardship by Native peoples of Wisconsin, USA. Ecol. Soc.

Watson, J.E.M., Dudley, N., Segan, D.B. & Hockings, M. (2014). The performance and potential of protected areas. Nature, 515, 67–73.

West, P., Igoe, J. & Brockington, D. (2006). Parks and Peoples: The Social Impact of Protected Areas. Annu. Rev. Anthropol., 35, 251–277.

Yibarbuk, D., Whitehead, P.J., Russell-Smith, J., Jackson, D., Godjuwa, C., Fisher, A., Cooke, P., Choquenot, D. & Bowman, D. (2001). Fire ecology and Aboriginal land management in central Arnhem Land, northern Australia: a tradition of ecosystem management. J. Biogeogr., 28, 325–343.

